# In silico tool for Predicting, Designing and Scanning IL-2 inducing peptides

**DOI:** 10.1101/2021.06.20.449146

**Authors:** Naman Kumar Mehta, Anjali Lathwal, Rajesh Kumar, Dilraj Kaur, Gajendra P. S. Raghava

## Abstract

Interleukin-2 (IL-2) based immunotherapy has been approved for treating certain types of cancer, as IL-2 plays a crucial role in regulating the immune system. In this study, we developed a method for predicting IL-2-inducing peptides. Our method was trained, tested, and validated on a main dataset containing 6,574 experimentally validated Major histocompatibility complex (MHC) binders, including 3,429 IL-2-inducing and 3,145 non-inducing peptides. A primary analysis of IL-2 inducing and non-inducing peptides revealed that certain residues, such as alanine and leucine, are more abundant in IL-2-inducing peptides. Initially, we developed alignment-based methods, which demonstrated high precision but limited coverage. Subsequently, we developed artificial intelligence-based models, including machine learning (ML), deep learning (DL), and large language models (LLM), to predict IL-2-inducing peptides. Our Extra Tree-based model, developed using dipeptide composition and peptide length, achieved a maximum AUC of 0.82. Finally, we constructed ensemble models that combined artificial intelligence and alignment-based methods. Our best ensemble model, which integrates the Extra Tree-based model with MERCI, achieved the highest AUC of 0.84 and an MCC of 0.51 on the main dataset. One limitation of the main dataset is that both IL-2-inducing and non-inducing peptides are MHC binders. To address this limitation, we created two additional datasets: Alternate Dataset 1, consisting of 3,429 IL-2-inducing peptides and 3,429 non-inducing peptides (MHC non-binders), and Alternate Dataset 2, consisting of 3,429 IL-2-inducing peptides and 3,439 non-inducing peptides (MHC binders + MHC non-binders). Our best ensemble model achieved AUCs of 0.9 and 0.8 with MCCs of 0.61 and 0.44 on Alternate Datasets 1 and 2, respectively. To assist the scientific community, we have integrated the best models from this study into a standalone software and web server, IL2pred, which enables users to predict, scan, and design IL-2-inducing peptides (https://webs.iiitd.edu.in/raghava/il2pred/).

**Highlights:**

- Development of alignment-based techniques for predicting IL-2-inducing peptides.
- Identification of amino acid patterns associated with IL-2 induction.
- Implementation of machine learning, deep learning, and LLM-based models for peptide prediction.
- Ensemble approaches combining alignment-based and AI-driven methods and integration into the webserver.

## 1. Introduction

Traditional cancer treatment strategies, such as chemotherapy and radiotherapy, have been the cornerstone of cancer care for decades. While effective, these approaches often come with significant short- and long-term side effects, including toxicity, fatigue, myalgias, cognitive dysfunction, and infertility[1]. Immunotherapy is often considered the “fifth pillar” of new-age therapies that aim to eliminate cancer cells via immune system activation. This can be achieved by targeting the tumor antigens or by passively enhancing the anti-tumor immune response by administering cultured lymphocytes [2]. Recent clinical studies have confirmed the efficacy and safety of immunotherapy in managing large cohorts of patients [3–5]. Several targeted immunotherapies (e.g., drug-target conjugate), therapeutic cancer vaccines (e.g. T-VEC), adoptive targeted therapies (e.g., CAR T-cells), and immunomodulators such as interleukins have been approved and marketed for use in different cancer types[2,6]. Interleukins are small glycoproteins that can have a pleiotropic effect on the immune system. They bind to receptors on the cell surface that can aid in immune cells’ differentiation, survival, and proliferation [7].

Currently, two interleukin-based drugs are approved by the Food and Drug Administration (FDA) for treating human malignancies. This includes the interferon-alpha-based drug Roferon-A and IL-2 based Aldesleukin or Proleukin [8]. IL-2 is an early cytokine that was commercially approved to treat metastatic renal carcinoma and metastatic melanoma by the FDA. IL-2 is a 15.5 KDa alpha-helical cytokine that belongs to the gamma chain family of immune modulators. CD4+ T-cells chiefly produce IL-2, but this can also be made by CD8+ T-cells, dendritic cells, and natural killer cells (NK-cells) [3]. While IL-2 is known for its immune-stimulating activity at low concentrations, it also promotes the expansion of regulatory T-cells and immune tolerance at higher concentrations, reflecting its dual role in immune regulation. IL-2 works through the JAK-STAT pathway and helps maintain and differentiate CD4+ T-cells in various subsets, including Th1, Th2, and Th17. It can promote CD8+ T-cell and NK-cell cytotoxic activity and the generation of memory cells [9]. The excellent immune-stimulating activity of IL-2 has led to the development of several treatment regimens. IL-2 has been used in clinical settings as a monotherapy agent and combined with several other therapeutic regimes. The low dose of IL-2 combined with interferon-alpha shows more significant therapeutic benefits for patients with renal cell carcinoma [10]. IL-2, when combined with chemotherapeutic agents, including cisplatin and dacarbazine, has been extensively investigated in patients with metastatic melanoma [11]. The IL-2, combined with targeted therapies such as gefitinib, showed a positive response in non-small cell lung cancer (NSCLC) patients [12]. Combining IL-2 with a therapeutic cancer vaccine also indicates a synergistic effect in advanced melanoma [13].

IL-2 therapy has a number of limitations, including a short half-life; high doses of IL-2 lead to toxicity, vascular leakage, and hypotension [14–16]. The clinicians and researchers employed several strategies for overcoming the challenges faced by IL-2 therapy. One such strategy widely employed by researchers in literature to improve the efficacy of IL-2 therapy is the generation of IL-2 inducing mutant peptides. The mutant version of IL-2 inducing peptides reportedly has low toxicity and can activate Natural-killer cells (NK-cells) without producing a high level of pro-inflammatory cytokines, thus preventing vascular leakage [7,17]. There are other mutants of IL-2 inducing peptides that can activate immune cells without pro-inflammatory activity and thus are free from toxicity-related problems [17]. These mutant versions of IL-2 inducing peptides are known as “Superkine’’ because of their enhanced antitumor property. These points highlight that the mutant version of natural IL-2 inducing peptides possesses a high therapeutic index. Thus, generating an improved version of IL-2 has become an important area of research. However, identifying and generating such mutants in clinical set-ups is time-consuming and cost-intensive. In-silico computation methods can help scientists and clinicians in this regard. Several past methods have also been developed that targeted and utilized the therapeutic potential of interleukin-based therapy for managing various human malignancies. CytoPred is one such method that is developed for the identification and classification of cytokines [18]. In addition, methods have been developed for inducing specific types of cytokines, including IL4pred [19], IL10Pred [20], IFNepitope [21], and IL6pred [22] for IL-4, IL-10, IFN-gamma, and IL-6, respectively. Despite the huge therapeutic potential, no method is available in the literature for identifying and generating peptides that can induce the natural secretion of IL-2. In the present study, a computational method for predicting and designing peptides that can potentially induce IL-2 cytokine has been developed.

## 2. Materials and Methods

### 2.1 Datasets

#### 2.1.1 Main dataset

A total of 8596 experimentally validated IL-2 inducing and non-inducing peptides were extracted from the largest repository of the immune epitope database (IEDB), filtering for MHC binders from any host organism that was experimentally confirmed to either induce or not induce IL-2 production. Of these, 4475 peptides were MHC binders, which can trigger IL-2 secretion as measured by different immunological assays. These epitopes were termed IL-2 inducing peptides and grouped under a positive set. We also extract 4121 MHC binding peptides that do not trigger the IL-2 secretion and are termed non-inducers. The MHC-binding peptides that do not induce IL-2 are called non-inducers and are grouped under a negative set. Literature evidence suggests that peptides of length between 8-25 are most suitable for MHC antigen processing and presentation. Thus, all peptides of length below 8 and above 25 were removed. Additionally, all the redundant peptides were removed. The final main dataset consists of 3429 IL-2 inducing and 3145 non-inducing peptides. One of the major features of our main dataset is all peptides are experimentally validated.

#### 2.1.2 Alternate Dataset 1

Our main dataset has all MHC binders, which means models developed on the main dataset are only suitable to MHC binders. In case the user does not know whether a given peptide is an MHC binder or a non-binder, then the model developed on the main dataset cannot be used. In order to overcome this limitation, we changed our negative dataset of non-induces from MHC binders to non-binders. In the Alternate dataset, we extracted and selected 3429 non-MHC-binding peptides from IEDB and assigned them as non-inducers. Finally, our Alternate dataset 1 contains 3429 MHC binding IL-2 inducing peptides as positive peptides and 3429 MHC non-binding IL-2 non-inducing peptides as negative peptides. Models developed on Alternate dataset 1 are suitable for predicting IL-2 inducing peptides in MHC non-binders.

#### 2.1.3 Alternate Dataset 2

Our models developed on our main dataset are suitable for predicting IL-2 inducing peptides in MHC binding peptides. Similarly, our models developed on Alternate dataset 1 are suitable to predict IL-2 inducing peptides in MHC non-binding peptides. In case the user has no idea whether their peptide is an MHC binder or a non-binder, then the above datasets are not suitable. In this study, we proposed alternate dataset 2 that contains 3429 MHC binding IL-2 inducing peptides, referred to as positive peptides and 3429 IL-2 non-induces. In Alternate Dataset 2, IL-2 non-inducers contain a mixture of MHC binders and non-binders, which do not induce IL-2.

### 2.2 Length distribution and composition analysis

To better understand the characteristics of the peptides in both the positive and negative datasets, a comprehensive analysis was performed on their length distribution and amino acid composition. This analysis was carried out using custom Python scripts, which were designed to generate bar plots for visual representation.

### 2.3 Positional preference analysis

In this analysis, a Two Sample Logo (TSL) was constructed to compare the proportional representation or relative abundance of amino acid residues between IL-2-inducing (positive) peptides and non-inducing (negative) peptides. The analysis was focused on identifying the preference for specific amino acids at particular positions within peptide sequences to distinguish between the two peptide groups. Notably, the first eight positions represent the N-terminal residues, while the final eight positions correspond to the C-terminal residues of the peptides.

### 2.4 Alignment-based approach

#### 2.4.1 Motif Search

Identification of the motifs within peptides is crucial in annotating their function. The MERCI software was used to identify specific motifs in both positive and negative datasets [23]. The software was implemented in two steps; firstly, the motifs for a positive dataset were extracted by providing the IL-2 inducing peptides as positive and non-inducing peptides as negative sets. In the second iteration, the motifs were retrieved for the negative dataset by inputting IL-2 non-inducing peptides as positive and IL-2-inducing peptides as negative sets. The motifs were calculated using the earlier approach in both positive and negative datasets. The different classification methods (NONE/BETTS-RUSSELL/KOOLMAN) were used to identify motifs in a mutually exclusive manner with the help of MERCI motif analyses. NONE applies no grouping; BETTS-RUSSELL categorizes amino acids into polar, hydrophobic, and small groups, while KOOLMAN focuses on aliphatic, aromatic, and other properties. The peptides containing unique motifs from both sets were screened to understand the overall coverage of the various motifs in the complete data.

#### 2.4.2 BLAST Search

This study utilised blastp-short to annotate IL-2 peptides based on their similarity to IL-2-inducing or non-inducing sequences [24]. Initially, a database was constructed using the “makeblastdb” command for IL-2 peptide sequences in the training dataset. The self-hits were excluded from the training dataset results and considered the top hit after removing them. The first hit was considered for the independent dataset to calculate results at various e-values.

### 2.5 AI-based classification method

#### 2.5.1 Feature Estimation

The different features of peptides present in both positive and negative datasets have been calculated. These calculated features are employed in developing ML-based prediction method development. For the feature generation, the Pfeature web server [25] was used, and the embeddings were generated from the pre-trained protBERT model [26]. With the help of Pfeature, 10,000 peptide descriptors were calculated for both positive and negative datasets. Pfeatutre includes I) Composition-based descriptors – Amino Acid Composition (AAC), Dipeptide Composition (DPC), Repetitive Residue Information (RRI), Physico-Chemical Properties (PCP), Distance Distribution of Residues (DDR); II) Binary profiles – Amino acid based binary profile (AAB); and LLM include Embeddings of Large Language Models trained on ProtBERT developed by Rostlab.

### 2.6 Prediction Models

#### 2.6.1 ML Models

Various ML-based algorithms have been implemented to classify IL-2 inducing and non-inducing peptides. The “scikit-learn” Python package was used for the classification. The classification algorithm includes - decision trees (DT), random forest (RF), multi-layer perceptron (MLP), eXtreme gradient boosting (XGBoost), support vector with the kernel as a radial basis (SVR), ExtraTreesClassifier (ET) and LASSO. Hyper-parameter tuning was implemented using the grid search CV technique with “ROC” as the optimization metric. The DT algorithm was based on the non-parametric supervised algorithm; RF is an ensemble-based method that fits numerous decision trees to predict the outcome of the dependent variable; KNN is an instance-based learning algorithm, and XGBoost is a tree-boosting classification algorithm based on an iterative search approach for making the final prediction.

#### 2.6.2 DL Models

For the DL approach, two distinct models were applied: first, a 1D CNN model tailored for processing sequential data and extracting hierarchical features along the sequence, and second, TabNet, a modern neural network architecture designed explicitly for tabular data [27]. The 1D CNN Model is particularly effective for sequential data processing, capturing patterns and dependencies along the peptide sequences. On the other hand, TabNet excels in tabular data by dynamically selecting and reasoning from features using sequential attention, making it suitable for our IL-2 inducing and non-inducing peptides dataset. Each model was fine-tuned independently on the datasets by adjusting hyperparameters to optimize their performance for the classification task.

#### 2.6.3 LLM Models

LLM are excellent at text classification because they deeply understand language. Trained on large amounts of text, they can be designed for specific tasks like classifying sequences, as was done in this research. LLMs are flexible and are particularly good at extracting important text features, allowing them to make accurate classifications even with limited data. This study used LLMs to differentiate between IL-2 and non-IL-2 sequences. The protBERT model was chosen and fine-tuned to match the characteristics of the datasets. After this fine-tuning, the model predicted whether each sequence belonged to the IL-2 or non-IL-2 category.

#### 2.6.4 Feature selection

As the literature suggests that not all descriptors are effective for developing machine learning models, three techniques were employed to select the best features: minimum Redundancy - Maximum Relevance (mRMR), SVC-L1 [28] and Recursive feature elimination (RFE). Feature selection was performed on DPC with length. The top 10, 100, and 200 features were computed using both methods on the datasets. After the feature selection step, all the features mentioned above were tested independently.

### 2.7 Ensemble method

In this approach, the best model obtained was combined with motif information from MERCI for reliable and biologically significant prediction of IL2-inducers. Along with the prediction models, the query peptide was searched against the test data with different thresholds for false positive (fp) values; if any motif was found in the peptide, the probability score of test data was increased by 0.5 for IL-2 inducing peptides and in the case of IL-2 non-inducers, the score was decreased by 0.5. This gives the probability of a sequence belonging to a specific class instead of a binary outcome.

### 2.8 Cross-validation

For developing the ML, DL, and LLM models, the standard protocols were followed as in previous studies [31498794]. Each dataset was divided into an 80:20 split for training and external validation, respectively. The main dataset contained 5,259 peptides for training and 1,315 peptides for testing. The alternate dataset 1 included 5,486 peptides for training and 1,372 for testing, while the alternate dataset 2 consisted of 5,487 peptides for training and 1,372 for testing. The different classifiers were trained and evaluated using a five-fold cross-validation method, a widely accepted technique for optimizing model parameters and performance. In the five-fold cross-validation, the training dataset was divided into five equal parts; iteratively, four parts were used for training and one for testing to fine-tune the model parameters. This process was repeated five times, ensuring each subset was used for both training and testing. All the classifiers were implemented using an in-house python script. Performance evaluation was based on metrics such as specificity, sensitivity, accuracy, MCC, and AUC. The statistical performance-evaluation parameters of the ML models are explained below:

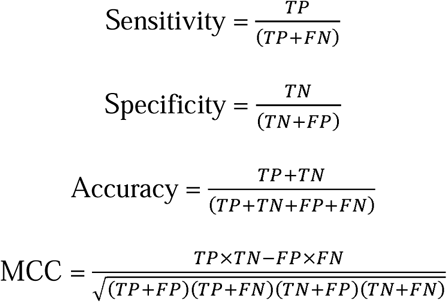

The overall workflow of the complete methodology is summarized below in **Figure 1**.

**Figure 1.**
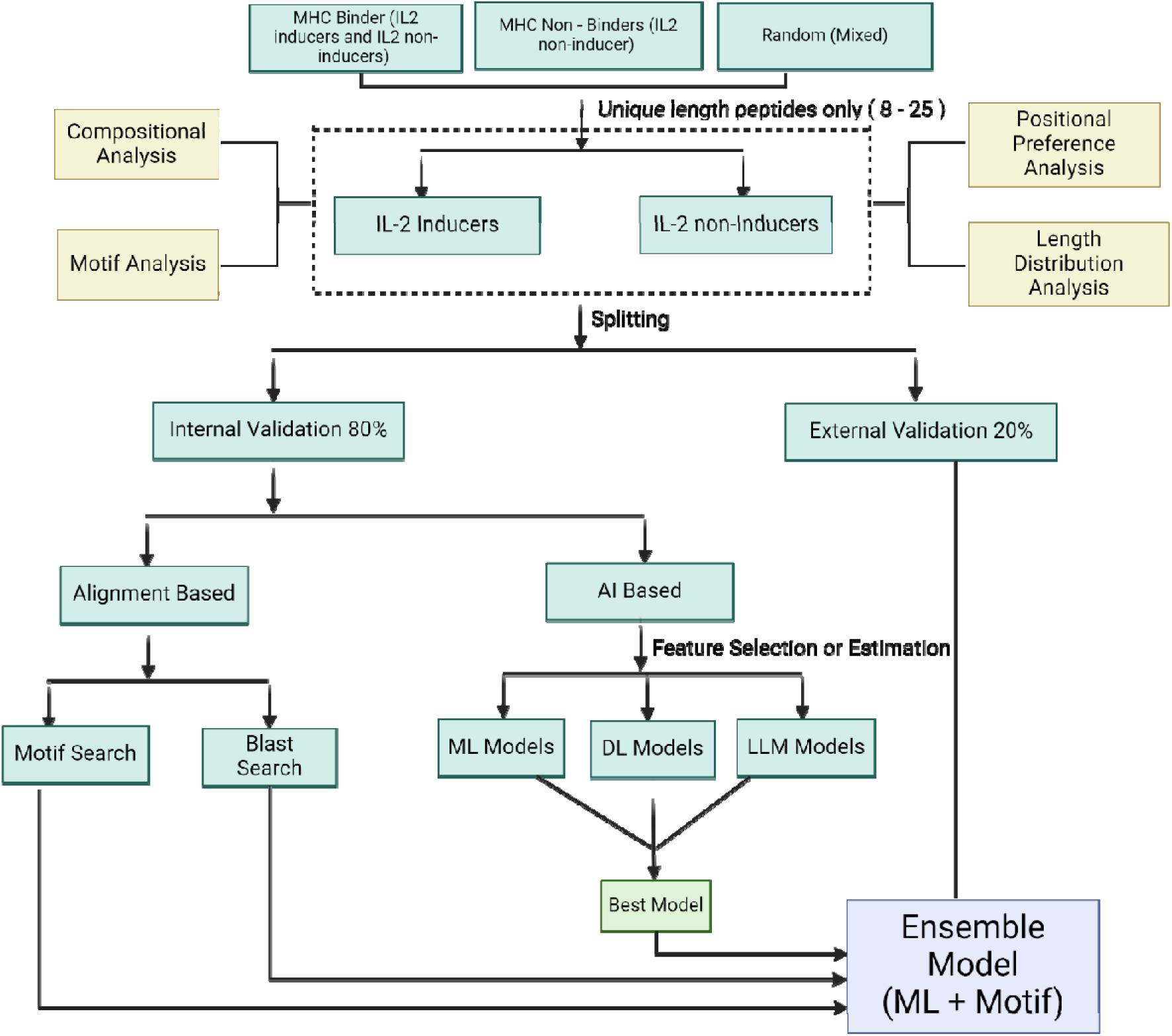
Schematic representation of the workflow of the study

### 2.8 Architecture of Web-server

A web server named “IL2pred” (https://webs.iiitd.edu.in/raghava/il2pred/) was developed to predict the IL-2 inducing and non-inducing peptides. The web server’s front end was developed using HTML5, JAVA, CSS3, and PHP scripts. It is based on responsive templates that adjust the screen based on the device’s size. It is compatible with almost all modern mobile, tablet, and desktop devices.

## 3. Results

In this study, three distinct datasets were used to develop the prediction algorithm – main dataset, alternate dataset 1 and alternate dataset 2. The distribution of peptides with respect to the MHC allele in our main dataset is available in Supplementary File 1 (S1.xlsx). All analyses and model building were conducted on these datasets. Based on this comprehensive analysis, different prediction models were built, trained, and externally validated to assess their performance in predicting IL-2 inducing and non-inducing peptides.

### 3.1 Dataset Length Distribution Analysis

Both positive and negative datasets were subjected to length distribution analysis. The length distribution analysis of both the positive and negative datasets of the main dataset is provided in Figure 2. The distribution analysis reveals that positive/IL-2 inducing peptides predominantly consist of peptide lengths ranging from 12-20, with a length of 15 as the most abundant. In negative/IL-2 non-inducing peptides, the peptides were mainly 15, 17, and 20-length amino acid sequences. The distribution analysis of the other two datasets has been reported in the Supplementary File 2: Figure SF1 and Figure SF2.

**Figure 2:**
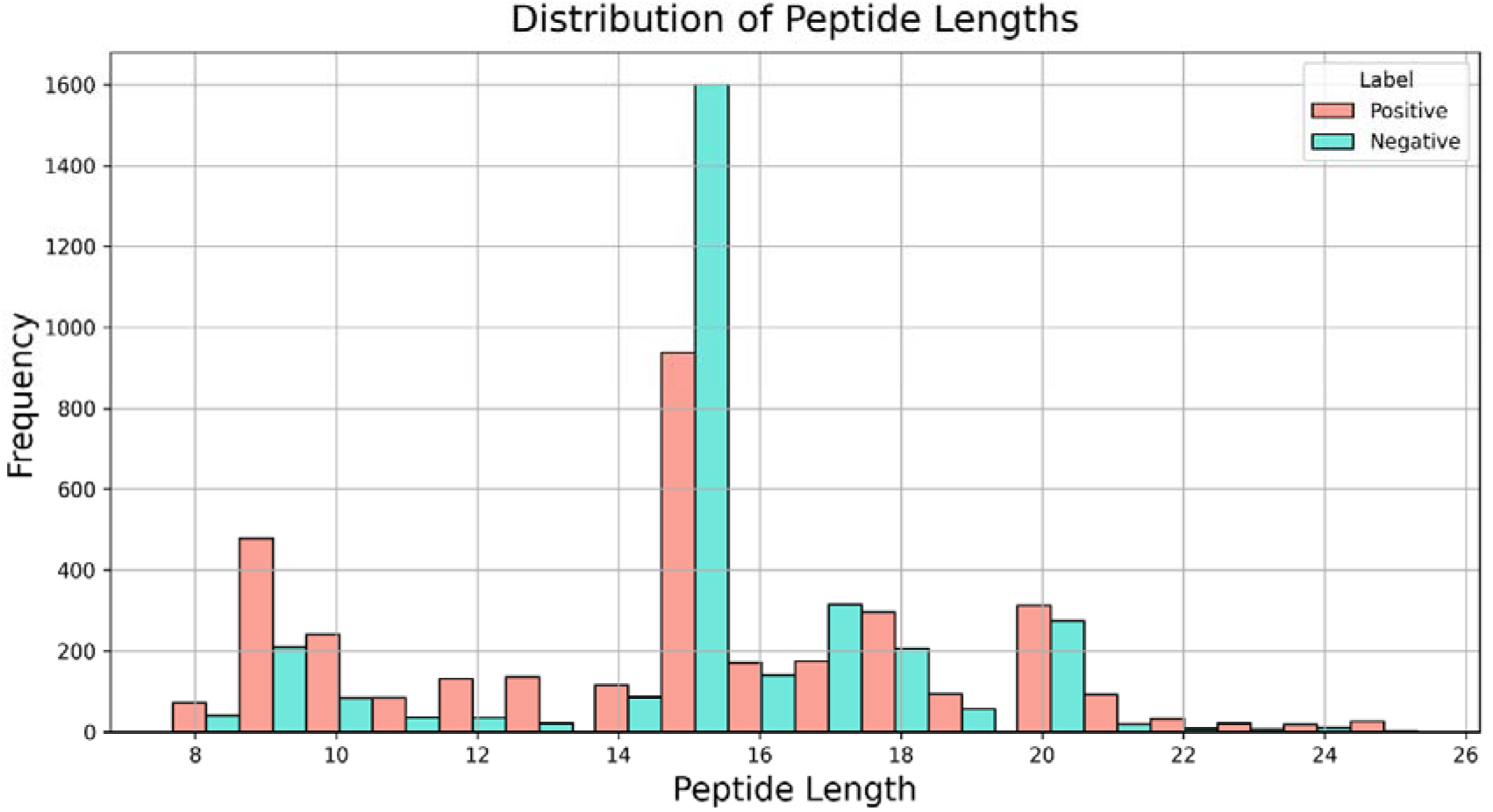
Histogram plot showing the frequency distribution of peptide length of the main dataset.

### 3.2 Average Amino Acid Composition Analysis

The positive and negative datasets were analysed for their average amino acid composition. In the main positive dataset, amino acid residues L, P, T, E, D, Y, C, A, Q, and F are highly significant (adjusted p-value < 0.05). The average amino acid composition analysis for the main dataset is provided in Figure 3, along with the adjusted p-value and the other two datasets are provided in Supplementary File 2: Figure SF3 and Figure SF4

**Figure 3:**
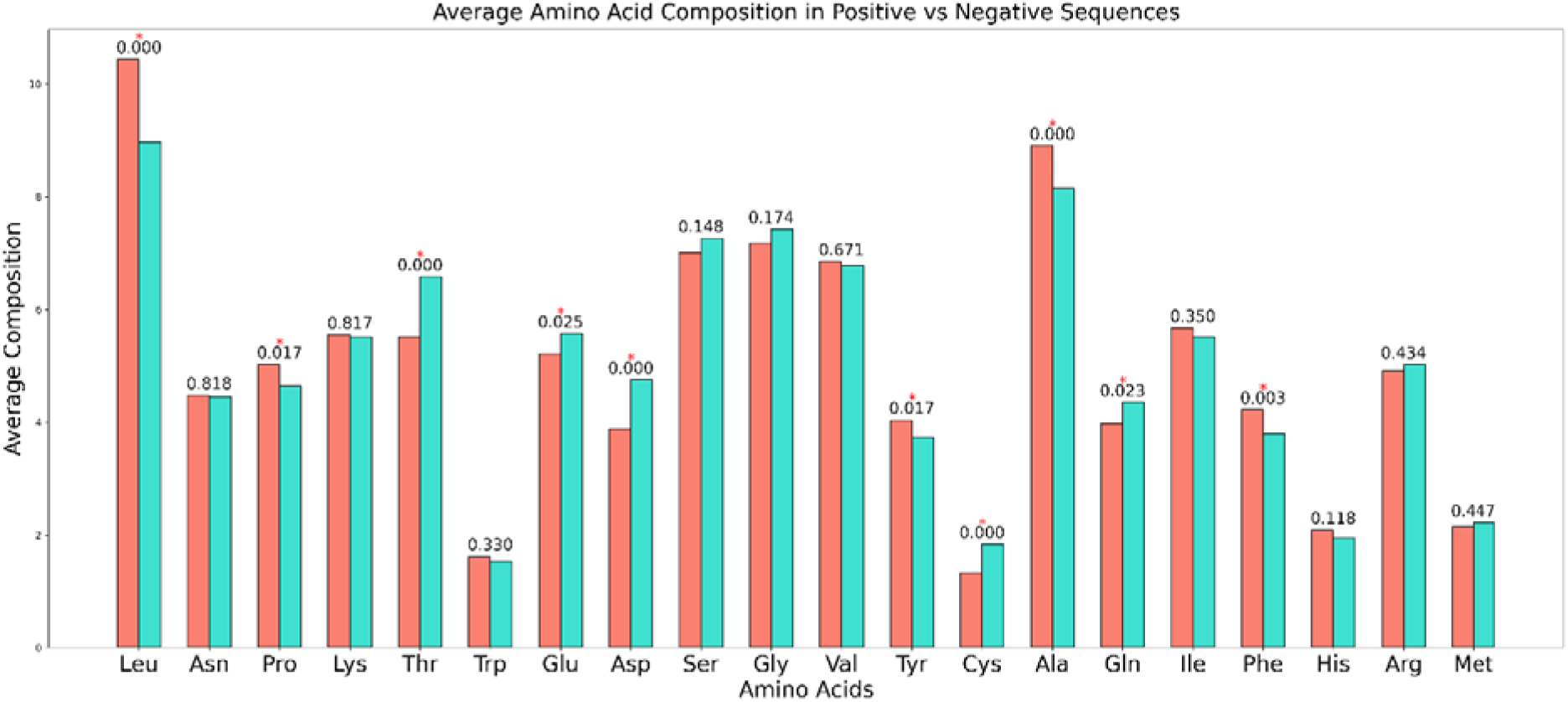
Bar plot of the peptides’ average single amino acid composition in the IL-2 inducers and non-inducers of the main dataset. The adjusted p-values are shown above the bars.

### 3.3 Positional preference analysis

The TSL reveals the positional preference of amino acids for positive and negative datasets. In the main positive dataset, IL-2 inducing peptides show a strong enrichment for hydrophobic residues like L, G, Y, and F and positively charged residues like R and K at specific positions. The depleted amino acids, such as T, D, S, and E, suggest that IL-2-inducing peptides avoid hydrophilic and acidic residues at these positions, as shown in Figure 4. The positional preference analysis for the other two datasets is provided in Supplementary File 2: Figure SF5 and Figure SF6.

**Figure 4:**
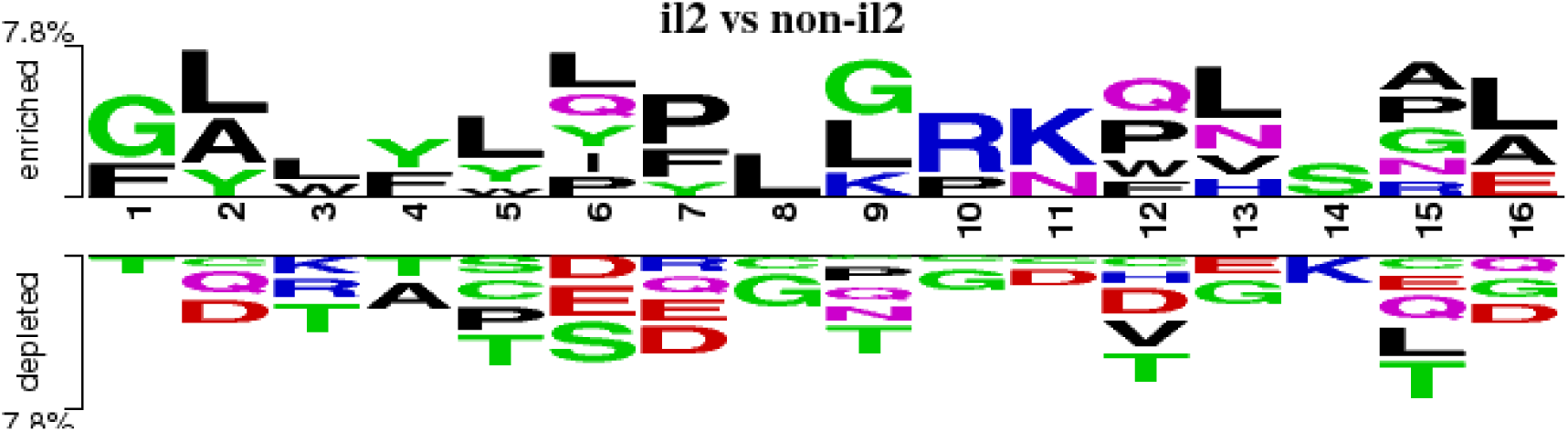
Two-sample logo displaying the positional conservation of amino acid for the main dataset.

### 3.4 Alignment-based approaches

#### 3.4.1 Motif Search

The MERCI software has been utilised to determine the motifs present exclusively in the IL-2 inducing peptides but not in IL-2 non-inducing peptides. Similarly, the motifs that are exclusive to IL-2 non-inducing peptides were computed. The different parameters and classification methods (NONE/BETS-RUSSELL/KOOLMAN) available in MERCI software have been utilised to differentiate the motifs into IL-2 inducing and non-inducing peptides. Among these, for the main dataset, the “NONE” classification scheme with fp 10 was selected as the best parameter because of its high accuracy despite having low coverage. The results of the top 10 motifs and the number of sequences from the independent datasets that occurred in the positive and negative classes of the main dataset are presented in Table 1. It is revealing that the motifs were hydrophobic in nature and also exclusive to IL-2 inducers. This result was also seen during the positional preference analysis. The results of motif coverage of the other two datasets were presented in Supplementary File 1 (S2.xlsx).

**Table 1:**
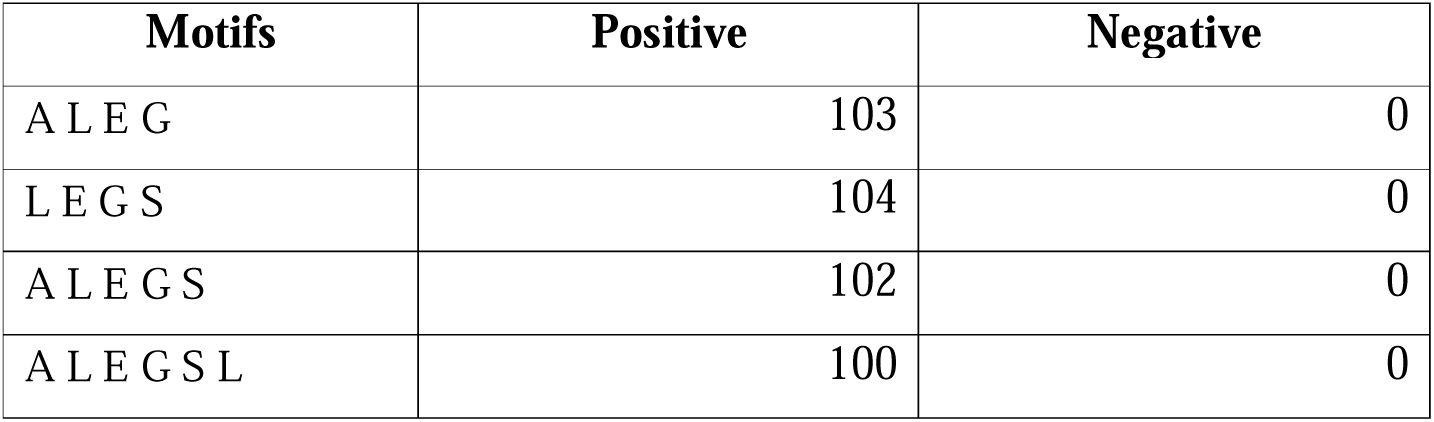
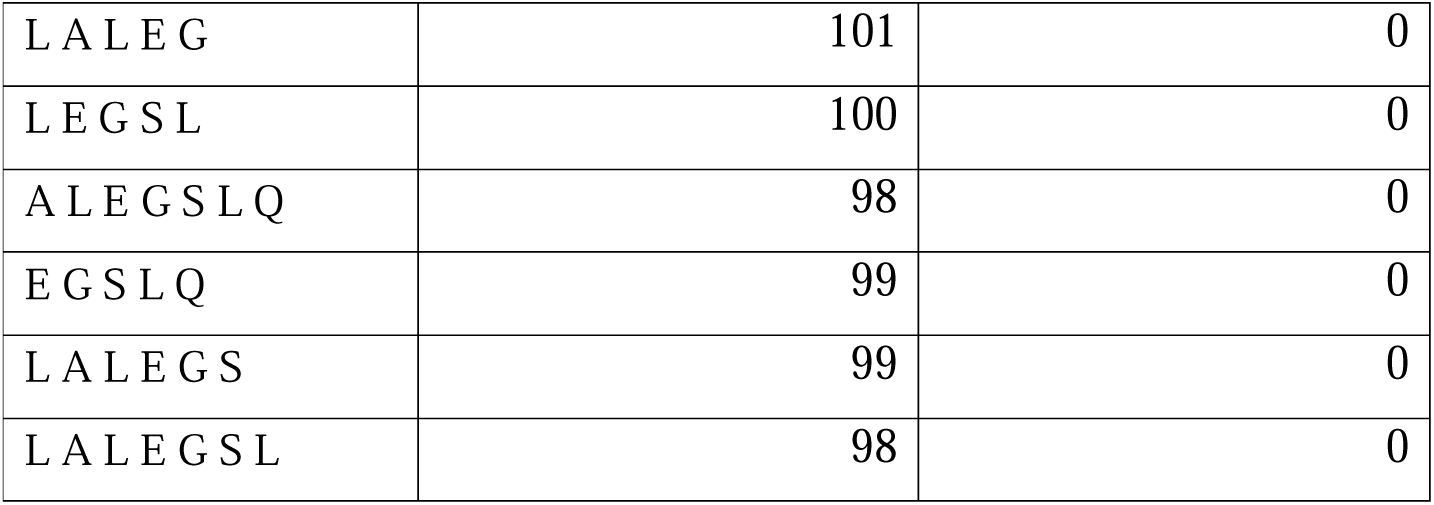
Top 10 IL-2 inducer motifs of the main dataset and the number of sequences in which they occurred.

| Motifs | Positive | Negative |
| --- | --- | --- |
| A L E G | 103 | 0 |
| L E G S | 104 | 0 |
| A L E G S | 102 | 0 |
| A L E G S L | 100 | 0 |
| L A L E G | 101 | 0 |
| L E G S L | 100 | 0 |
| A L E G S L Q | 98 | 0 |
| E G S L Q | 99 | 0 |
| L A L E G S | 99 | 0 |
| L A L E G S L | 98 | 0 |

#### 3.4.2 BLAST Search

The blastp-short was applied to perform a similarity search against a training dataset (main dataset) of IL-2 inducing and IL-2 non-inducing peptides. The search used blastp-short with an e-value range from 10^-7^ to 10^-1^ (Table 2). Blastp-short achieved optimal performance at an e-value of 10^-3^, correctly identifying 2100 IL-2 induced peptides and 402 IL-2 non-induced peptides, with 272 incorrect hits. E-values lower than 10^-3^ did not provide adequate sequence coverage, while values higher than 10^-3^ exhibited a higher error rate. The complete results for BLAST from e-values 10^-7^ to 10^-1^ for the other two datasets are detailed in the Supplementary File 1 (S3.xlsx).

**Table 2:** BLAST results for test datasets searched against the training datasets (main dataset).

| **E-value** | **Total hits** | **Correct positive hits** | **Incorrect positive hits** | **Correct negative hits** | **Incorrect negative hits** |
| --- | --- | --- | --- | --- | --- |
| 0.1 | 4644 | 2790 | 334 | 1178 | 342 |
| 0.01 | 3584 | 2374 | 225 | 762 | 223 |
| 0.001 | 2774 | 2100 | 121 | 402 | 151 |
| 0.0001 | 2164 | 1812 | 581 | 191 | 103 |
| 0.00001 | 901 | 743 | 31 | 67 | 60 |
| 0.000001 | 226 | 174 | 7 | 17 | 28 |

### 3.5 AI-based classification methods

#### 3.5.1 ML models

Various ML-based prediction models were trained, including DT, RF, KNN, MLP, ET, SVR, XGB and Lasso. First, the features of IL-2 and IL-2 non-inducing peptides using Pfeature were computed. Using only DPC as the feature, the ET method yielded the best results with an AUC of 0.81 and an MCC of 0.45 on the test data of the main dataset. Length was used as an additional feature to recognise the importance of peptide length in MHC antigen presentation and IL-2 induction. The ET-based model using these hybrid features (DPC and peptide length) achieved a higher AUC of 0.82 and an MCC of 0.48 on the test set. The details for the best classifiers against each feature set for the independent dataset of the main dataset have been provided in Table 3. The statistical details for different classifiers using various feature sets for all three datasets are in the Supplementary File 1 (S4, S5 and S6.xlsx).

**Table 3:**
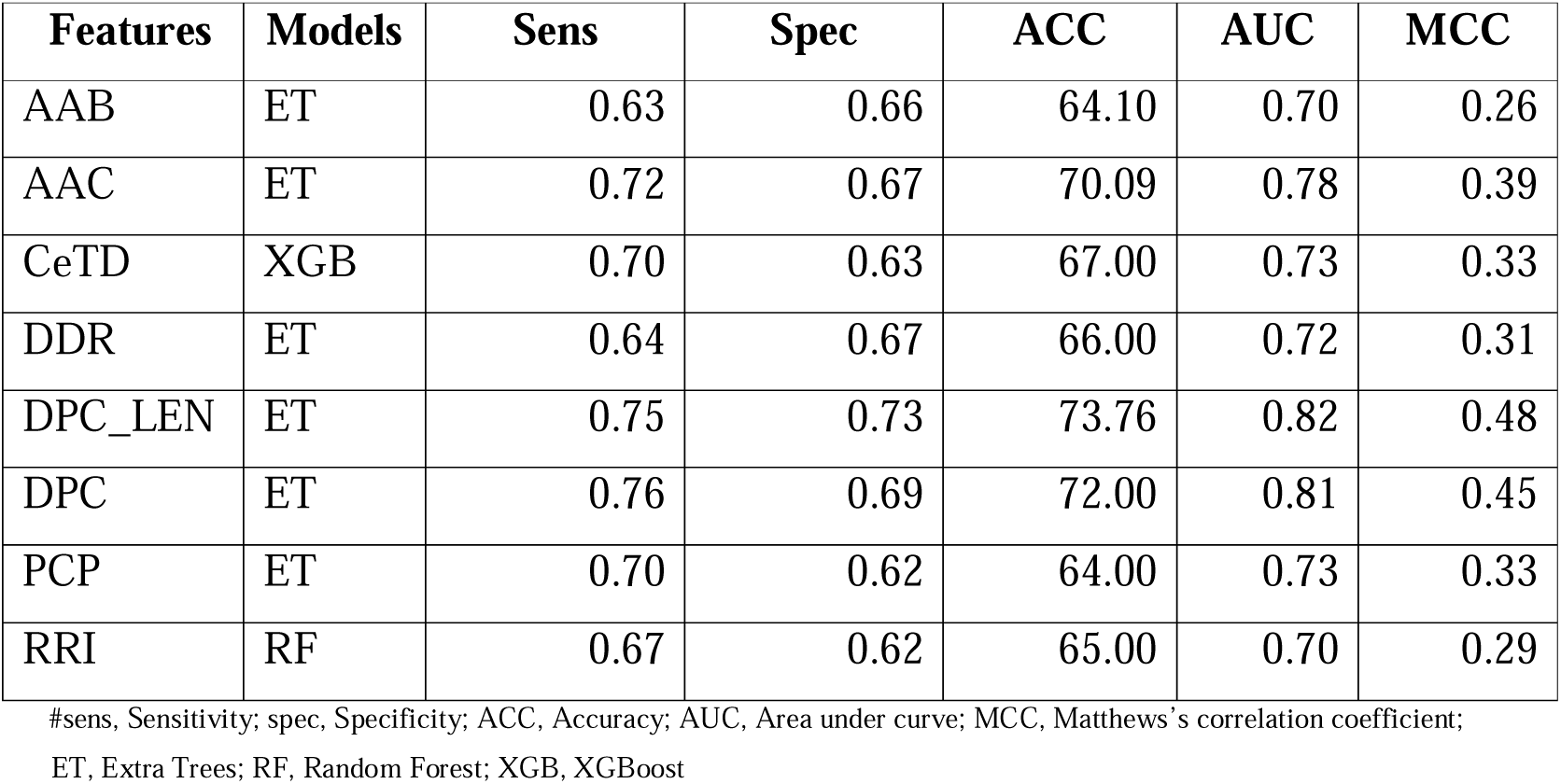
The test data performance for the best classifiers developed using various types of peptide features for the main dataset (only the best models are shown):

#### 3.5.2 DL models

Using the TabNet classification model, a maximum AUC of 0.65 and an MCC of 0.20 was achieved on the test data of the main dataset with Compositional Enhance Transition and Distribution (CeTD) as a feature.

For the 1D CNN model, an AUC of 0.71 and an MCC of 0.32 were achieved on the test data of the main dataset using DPC-length features. The top 3 best performances for the main dataset, using the independent dataset, have been provided in Table 4 and the detailed results for the TabNet and CNN models applied to IL-2 sequences for the other two datasets are provided in Supplementary File 1 (S7.xlsx).

**Table 4:**
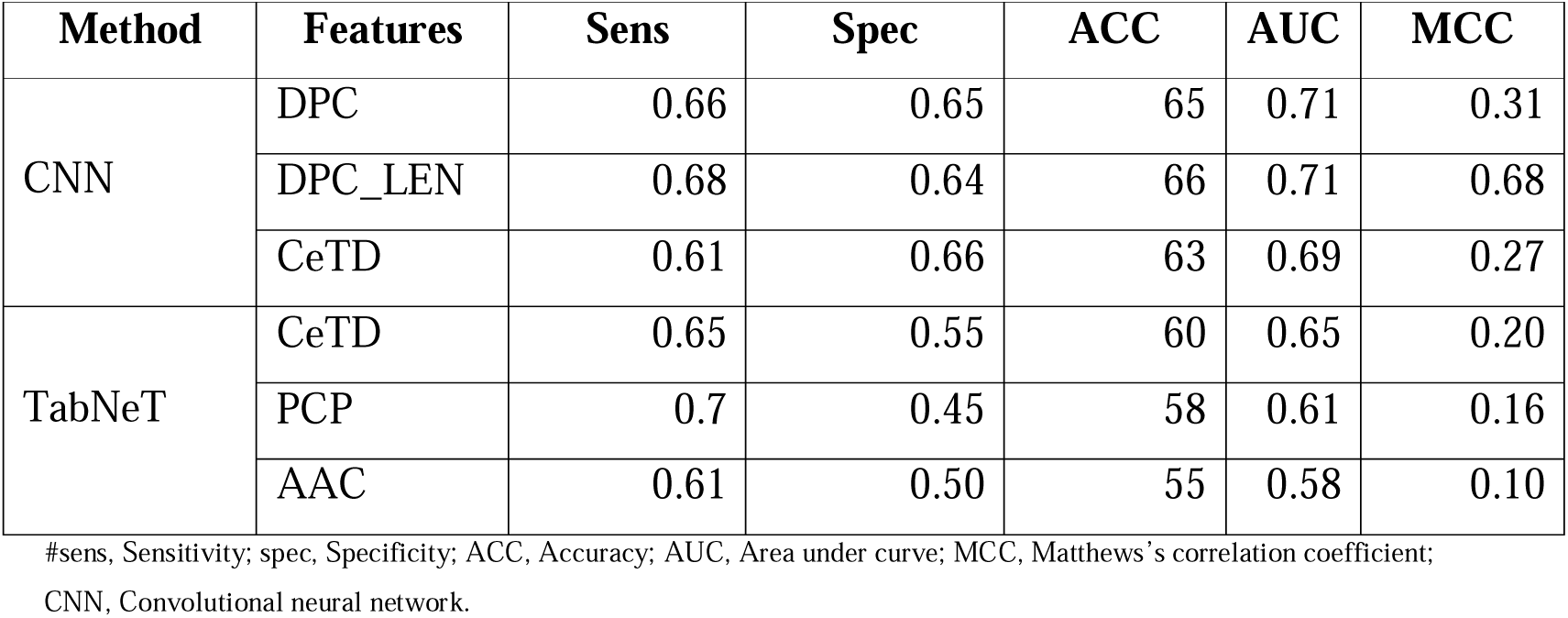
The top 3 best feature performances of the DL in the main dataset:

| Method | Features | Sens | Spec | ACC | AUC | MCC |
| --- | --- | --- | --- | --- | --- | --- |
| CNN | DPC | 0.66 | 0.65 | 65 | 0.71 | 0.31 |
|  | DPC_LEN | 0.68 | 0.64 | 66 | 0.71 | 0.68 |
|  | CeTD | 0.61 | 0.66 | 63 | 0.69 | 0.27 |
| TabNeT | CeTD | 0.65 | 0.55 | 60 | 0.65 | 0.20 |
|  | PCP | 0.7 | 0.45 | 58 | 0.61 | 0.16 |
|  | AAC | 0.61 | 0.50 | 55 | 0.58 | 0.10 |

#sens, Sensitivity; spec, Specificity; ACC, Accuracy; AUC, Area under curve; MCC, Matthews's correlation coefficient; CNN, Convolutional neural network.

#### 3.5.3 LLM models

For this study, a pre-trained large language protBERT model was utilised. The model was fine-tuned for specific tasks by adjusting the number of epochs. Among the epochs tested, epoch 5 provided the best results, with a maximum AUC of 0.69 and an MCC of 0.21 for the main dataset.

Furthermore, since embeddings generated by the fine-tuned model can serve as valuable features, these features were extracted and applied in various ML algorithms. The ET method was the best-performing model, achieving an AUC of 0.71 and an MCC of 0.31. The performance of the Fine-tuned and best classifier model for the main dataset has been provided in Table 5, and the details results for this LLM model for each dataset are given in Supplementary File 1. (S8.xlsx)

**Table 5:** The performance of the Fine-tuned and best classifier model for the main dataset:

| Epochs | Feature | Model | Sens | Spec | ACC | AUC | MCC |
| --- | --- | --- | --- | --- | --- | --- | --- |
| **3** | Sequence | Fine-tuned | 0.62 | 0.68 | 65.25 | 0.68 | 0.30 |
|  | Embeddings | ET | 0.64 | 0.68 | 66.00 | 0.71 | 0.32 |
| **5** | Sequence | Fine-tuned | 0.82 | 0.37 | 60.61 | 0.69 | 0.21 |
|  | Embeddings | RF | 0.69 | 0.62 | 65.00 | 0.71 | 0.31 |
| **7** | Sequence | Fine-tuned | 0.90 | 0.26 | 60.23 | 0.60 | 0.23 |
|  | Embeddings | ET | 0.74 | 0.60 | 67.00 | 0.72 | 0.34 |

#sens, Sensitivity; spec, Specificity; ACC, Accuracy; AUC, Area under curve; MCC, Matthews's correlation coefficient;
ET, Extra Trees; RF, Random Forest.

#### 3.5.4 Feature Selection

Three feature selection methods – mrMR, SVC-L1, and RFE were applied to the DPC-length feature on each dataset. The best model’s AUC for the main dataset decreased with all three methods when 10, 50 and 100 features were selected. When 200 features were selected, the model’s AUC remained at approximately 0.82 for each method. The Performance metrics results of our feature selection approach on the main dataset using SVC-L1, mRMR, and RFE have been provided in Table 6. The selected features were reported in the Supplementary File 1 (S9.xlsx).

**Table 6:** Performance metrics results of our feature selection approach on the main dataset using SVC-L1, mRMR and RFE:

| Type | Features | Model | Number | Sens | Spec | ACC | AUC | MCC |
| --- | --- | --- | --- | --- | --- | --- | --- | --- |
| mRMR | DPC_LEN | RF | 10 | 0.68 | 0.68 | 65 | 0.69 | 0.31 |
|  |  | ET | 100 | 0.70 | 0.67 | 69 | 0.76 | 0.37 |
|  |  | ET | 200 | 0.74 | 0.71 | 73 | 0.81 | 0.45 |
| SVC-L1 | DPC_LEN | ET | 10 | 0.68 | 0.68 | 65 | 0.70 | 0.30 |
|  |  | RF | 100 | 0.71 | 0.69 | 70 | 0.77 | 0.40 |
|  |  | RF | 200 | 0.71 | 0.75 | 73 | 0.81 | 0.46 |
| RFE | DPC_LEN | ET | 10 | 0.58 | 0.72 | 65 | 0.69 | 0.31 |
|  |  | RF | 100 | 0.72 | 0.70 | 71 | 0.78 | 0.43 |
|  |  | ET | 200 | 0.76 | 0.74 | 75 | 0.82 | 0.50 |

#sens, Sensitivity; spec, Specificity; ACC, Accuracy; AUC, Area under curve; MCC, Matthews's correlation coefficient.

### 3.6 Ensemble method

To enhance the predictive capability of our model, we employed an ensemble approach in this study. The highest AUC and MCC were achieved using the ensemble method compared to all other methods, including ML, DL, and LLM on the main dataset. Combining our top-performing model, DPC-length, with ET, along with motif scores using fp set to 10 and “NONE” as the classification method, an AUC of 0.84 and an MCC of 0.51 was obtained. We have tried other classification methods also, but this approach surpassed all other methods evaluated, highlighting its effectiveness in predicting biologically significant IL-2 inducers. The performance of the top 10 motifs on fp 10 developed using hybrid features (DPC-LEN and motif score) on all three datasets can be found in Table 7. The statistical details of various classifiers built using the hybrid feature set (DPC+Length) have been presented in Supplementary File 1 for all the datasets (S10, S11 and S12.xlsx).

**Table 7:** The performance of the top 10 motifs on fp 10 developed using hybrid features (DPC-LEN and motif score) on all three datasets.

| Dataset | Classification | Sens | Spec | ACC | AUC | MCC |
| --- | --- | --- | --- | --- | --- | --- |
| Main dataset | None | 0.77 | 0.74 | 76 | 0.84 | 0.51 |
|  | KOOLMAN | 0.77 | 0.73 | 75 | 0.83 | 0.50 |
|  | BETTS-RUSSELL | 0.76 | 0.73 | 75 | 0.82 | 0.50 |
| Alternate dataset 1 | None | 0.77 | 0.84 | 80 | 0.90 | 0.61 |
|  | KOOLMAN | 0.75 | 0.83 | 79 | 0.87 | 0.59 |
|  | BETTS-RUSSELL | 0.77 | 0.83 | 80 | 0.88 | 0.60 |
| Alternate dataset 2 | None | 0.71 | 0.73 | 73 | 0.80 | 0.44 |
|  | KOOLMAN | 0.70 | 0.73 | 72 | 0.79 | 0.43 |
|  | BETTS-RUSSELL | 0.69 | 0.75 | 72 | 0.79 | 0.44 |

#sens, Sensitivity; spec, Specificity; ACC, Accuracy; AUC, Area under curve; MCC, Matthews’s correlation coefficient.

## 4. Modules and Functionalities of IL2pred Web-server

This web server is compatible with all sorts of devices, viz. Desktop, tablets, and phones and hence provide an interactive and better experience to the users. The web-server has four main modules – 1) Prediction; 2) Design; 3) Protein Scan; 4) Motif Scan. The “Prediction module” allows users to identify the peptides with IL-2 inducing potential. Users of the web server can submit the peptides of interest in FASTA format as well as upload the FASTA format files from their local machine. The results of the user-defined query are displayed on the HTML page as IL2-inducer and IL-2-non-inducer, along with the prediction score and selected properties. The results of the user-defined query can also be downloaded from the web server in the CSV format. The “Design module” is of utmost importance for a biologist who wishes to design new, potent peptides with enhanced IL-2 inducing potential. The input for the design module is a single-line peptide sequence file, which can either be pasted in the box provided on the module page or locally uploaded onto the web server site. Once the user-defined query is submitted on the web server, the algorithm implemented on the “Design Module” generates all possible mutants of the given query peptide sequence and then ranks them based on their IL-2 inducing potential. The ranking algorithm is of greater importance as it ranks the newly generated single-letter mutant peptides for their ability to induce IL-2. Thus, it helps biologists narrow down their choice of selecting the best mutant peptide with enhanced IL-2 inducing ability. The prediction score and the other predicted metrics are displayed on the web server site, which can be downloaded as a CSV file for further use. The “Protein Scan” module lets users predict IL-2 inducing regions within a given protein sequence. Users can submit the protein sequence of their interest in a single-line format. The algorithm generates all possible fragments of the user-defined length and then predicts the IL-2 inducing potential of all generated fragments overlappingly. This will help in the identification of IL-2 rich and scarce regions in a given protein sequence and thus help biologists in selecting and prioritizing appropriate regions of protein for their study the last model is the Motif scan, which will allow users to find crucial motifs present in the query protein sequence to find out its role as an IL-2 inducer or IL-2 non-inducer. The results from this module can be downloaded in CSV format. The user-friendly web server, along with all functionalities, is available at (https://webs.iiitd.edu.in/raghava/il2pred).

## 5. Discussion

The high-throughput studies have significantly enriched our knowledge regarding the tumor onset to progression. However, the survival of patients is dismal as a whole, and finding novel therapeutics remains a significant challenge. Evidence from pre-clinical and clinical studies advocates activating the immune system to treat human malignancies [7]. Recently, various peptide-based prediction models have been developed using machine learning approaches methods such as random forest, support vector machine, deep neural networks, and several others that are instrumental in paving advanced ways for successful peptide-based therapeutics, diagnostics and drug discovery [29]. Some such methods are the identification of IL-6, IL-10 & IL-4 inducing peptides [19,20,22]; others predict the anti-microbial [30][29374199], anti-cancer [31], and anti-/pro-inflammatory peptides [32,33], using different properties such as distribution patterns of amino acids, structure-based properties generated from the peptide sequences. Pro-inflammatory cytokines are mediators of various biological processes controlled by the body. Uncontrolled levels of these cytokines can lead to many pathological conditions, such as cancer and autoimmunity. Recently, recombinant proteins have been used as biological drugs that can help neutralize an inflammatory cytokine or block its receptors. Interleukin-1 (IL-1) was used to treat anaemic patients and the ones undergoing chemotherapy. However, being a potent proinflammatory cytokine, it resulted in significant side effects, namely fever, headache, fatigue, etc [34]. Cytokine therapy has emerged to increase anti-tumor immunity using lymphocyte activators such as IL-2. IL-2 is critical to the survival, growth, and activation of T-cells and NK cells; it showed clear anti-tumor effects. Unfortunately, it had severe side effects associated with it, such as vesicular leakage, edema, and hypotension. However, considering its importance in immunotherapy, novel mutants of IL-2, such as FSD13, were designed to handle the side effects to a large extent and showcased improved anti-tumor effects with limited toxicity [35]. As mentioned above, IL-2 is one such cytokine that can boost the antitumor immune response by maintaining and promoting the activity of CD4+ T-cells and cytotoxic CD8+ T-cells. Thus, identifying peptides or antigenic regions that can boost the production of IL-2 would become an important consideration while designing cancer treatment regimens and vaccine candidates.

The present study is a systematic attempt to make an in-silico model for predicting the IL-2 inducing capability of the peptide/antigenic region. The in-silico model is developed on the sequence features of IL-2 inducing and non-IL-2 inducing peptides extracted from the largest repository of experimentally validated immune epitopes database. The non-redundant dataset ensures that no overfitting/bias/noise should affect the developed model due to the presence of multiple instances of the same peptide. Literature evidence suggests that the length of the peptide can affect its binding to the MHC complex and, thus, immune activation [36]. The length distribution analysis reveals that IL-2 inducing peptides predominantly consist of lengths ranging from 12-20, while for non-IL-2 inducers, it varies from 15-17. The compositional analysis reveals that IL-2 inducing peptides are enriched in A, L, S, and Y residues. Past studies also reveal that peptides enriched in S, L, and A residues have increased anti-tumor capabilities [37] as they can induce apoptosis in cancer cells via modulation of autophagy [38]. Thus, we can suggest that sequence composition can also be used to discriminate IL-2 inducing peptides from the non-IL-2 inducing peptides to some extent.

In addition to the above, the development of the IL-2 prediction models was also carried out using several other sequence features on multiple machine learning-based algorithms. It is observed that the DPC-based ET-model can discriminate between the IL-2 inducing and non-IL-2 inducing peptides with good accuracy. However, the hybrid model developed on DPC and length of peptides with motif score performed better than the DPC-length model alone in terms of balanced model performance measures. Both models were incorporated into the web server. We believe the developed tool will help the scientific community better understand IL-2 inducing/non-IL-2 inducing peptides. However, with the increased number of organism-specific datasets, the applicability of the developed model can further be improved. We anticipate that the developed tool may be of great use to the scientific community for identifying IL-2 inducing/non-IL-2 inducing peptides from the proteomes for improving the current cancer immunotherapeutics. Also, the developed tools can generate the mutant peptides that are predicted to have IL-2 inducing potential.

## 6. Conclusion

Past studies strongly reveal that direct administration of IL-2 may have several toxic effects. The capability of an antigenic region or mutant peptide to induce the IL-2 response is of great significance for activating the immune response towards malignant tissue. However, identifying such mutant IL-2 inducing peptides with conventional experimental techniques is time-consuming and costly. Therefore, a computational tool in the form of a web server is provided to the scientific community for predicting and identifying the IL-2 inducing/non-inducing peptides from the given proteomes. The web server is freely available to the scientific community at https://webs.iiitd.edu.in/raghava/il2pred/. We hypothesize that the developed tools will be highly utilized by the scientific community for identifying and prioritizing potential immunotherapeutic candidates.

## Author’s Contribution

**Naman Kumar Mehta:** Webserver implementation, Software and Coding, Investigation, Data curation, writing original draft, Reviewing and editing the original draft**. Anjali Lathwal:** Conceptualization, Methodology, Validation, Formal analysis, Webserver implementation, Software and Coding, Investigation, Data curation, writing original draft, Reviewing and editing the original draft. **Rajesh Kumar**: collected and processed the datasets. **Dilraj Kaur**: Webserver implementation, Reviewing and editing the original draft. **Gajendra Pal Singh Raghava**: Supervision, Conceptualization, project administration, Reviewing and editing original draft, Investigation, Validation.

## Supplementary Materials

### Supplementary File 1

There are 12 different sheets in this Excel file: (1) The distribution of peptides with respect to the MHC allele in our main dataset (2) The results of motif analysis of alternate dataset 1 and 2 (3) The complete results for BLAST from e-values 10^-7^ to 10^-1^ for the alternate dataset 1 and 2 (4) The statistical details for different classifiers using various feature sets for main dataset (5) The statistical details for different classifiers using various feature sets for alternate dataset 1 (6) The statistical details for different classifiers using various feature sets for alternate dataset 2 (7) The detailed results for the TabNet and CNN models applied to IL-2 sequences for each dataset (8) The results for this LLM model for each dataset. (9) List of top 200 feature names for each dataset computed using RFE. (10) The statistical details of various classifiers built using the hybrid feature set (DPC+Length) for the main dataset. (11) The statistical details of various classifiers built using the hybrid feature set (DPC+Length) for the alternate dataset 1. (12) The statistical details of various classifiers built using the hybrid feature set (DPC+Length) for the alternate dataset 2.

### Supplementary File 2

The supplementary file contains four figures: **Figure SF1.** Histogram plot showing the frequency distribution of peptide length of the alternate dataset 1. **Figure SF2.** Histogram plot showing the frequency distribution of peptide length of the alternate dataset 2. **Figure SF3.** Bar plot of the peptides’ average single amino acid composition in the IL-2 inducers and non-inducers of the alternate dataset 1. The adjusted p-values are shown above the bars. **Figure SF4.** Bar plot of the peptides’ average single amino acid composition in the IL-2 inducers and non-inducers of the alternate dataset 2. The adjusted p-values are shown above the bars. **Figure SF5.** Two-sample logo displaying the positional conservation of amino acid for the alternate dataset 1. **Figure SF6.** Two-sample logo displaying the positional conservation of amino acid for the alternate dataset 2.

## Data Availability Statement

The datasets generated for this study can be accessed on the ’il2pred’ web server at https://github.com/raghavagps/il2pred. The source code for this study is publicly available on GitHub and can be found at https://github.com/raghavagps/il2pred.

## Acknowledgement

The authors express their gratitude to the University Grants Commission (UGC), Council of Scientific and Industrial Research (CSIR), and Department of Science & Technology (DST) for their generous fellowships and financial support. They also thank the Department of Computational Biology, IIITD, New Delhi, for the excellent infrastructure and facilities. The authors would like to acknowledge the Department of Biotechnology (DBT) for the infrastructure grant awarded to the institute. Furthermore, they would like to acknowledge BioRender.com for the creation of the figures utilized in this work.

## Author’s Biography

1. Naman Kumar Mehta is currently pursuing a PhD in Computational Biology at the Department of Computational Biology, Indraprastha Institute of Information Technology, New Delhi, India.
2. Anjali Lathwal is currently working as an assistant professor in the Department of Computer Science and Engineering at IGDTUW Delhi.
3. Rajesh Kumar is currently working at the National Cancer Institute (NCI), Bethesda, Maryland, USA.
4. Dilraj Kaur is currently working at the Centre of Cancer Drug Discovery, Institute of Cancer Research (ICR), London, UK.
5. Gajendra P. S. Raghava is currently working as a Professor and Head of the Department of Computational Biology, Indraprastha Institute of Information Technology, New Delhi, India.

## List of Abbreviations

IL-2: Interleukin 2
MHC: Major Histocompatibility Complex
FDA: Food and Drug Administration
IEDB: Immune Epitope Database
AAB: Amino Acid Binary Profile
AAC: Amino Acid Composition
CeTD: Composition Enhanced Transition Distribution
DDR: Distance Distribution of Residues
DPC_LEN: Dipeptide Composition with length
PCP: Physico-Chemical Properties Composition
RRI: Repetitive Residue Information

## Key Points

- IL-2 is an immunoregulatory cytokine that plays a key role in cell-mediated immunity.
- IL-2 has been approved by the FDA as a therapeutic cytokine for the treatment of various cancer types, either in combination or in mutated form.
- Conventional experimental approaches pose a problem in identifying, prioritizing, and designing novel IL-2 peptides or proteins, thus limiting their clinical utility.
- IL2pred is a highly accurate in-silico method developed for predicting the IL-2 inducing peptides/proteins.
- IL2pred algorithm is implemented in the form of a webserver for the usage of the scientific community.

