## Supplementary File 2 for "In silico tool for Predicting, Designing and Scanning IL-2 inducing peptides"

**Supplementary Information**

**Figure SF1:** Histogram plot showing the frequency distribution of peptide length of the alternate dataset 1.

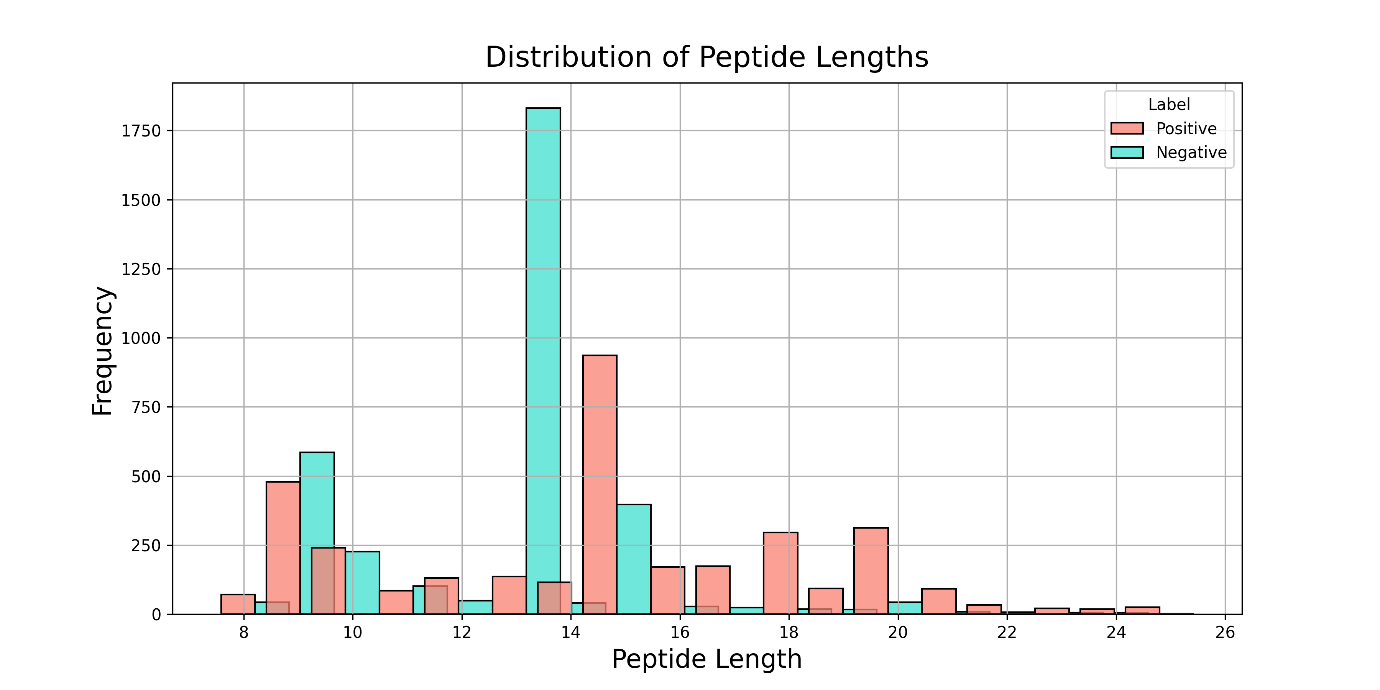

**Figure SF2:** Histogram plot showing the frequency distribution of peptide length of the alternate dataset 2.

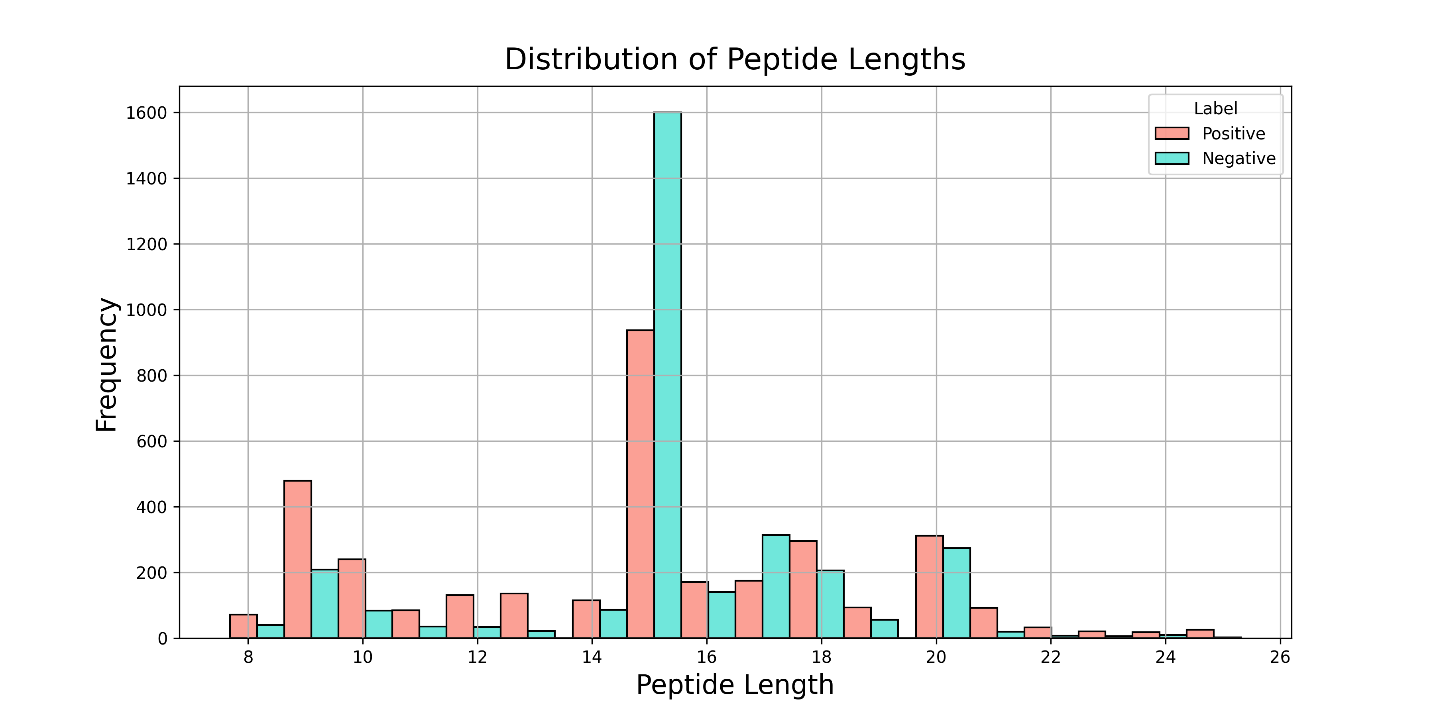

**Figure SF3:** Bar plot of the peptides' average single amino acid composition in the IL-2 inducers and non-inducers of the alternate dataset 1. The adjusted p-values are shown above the bars.

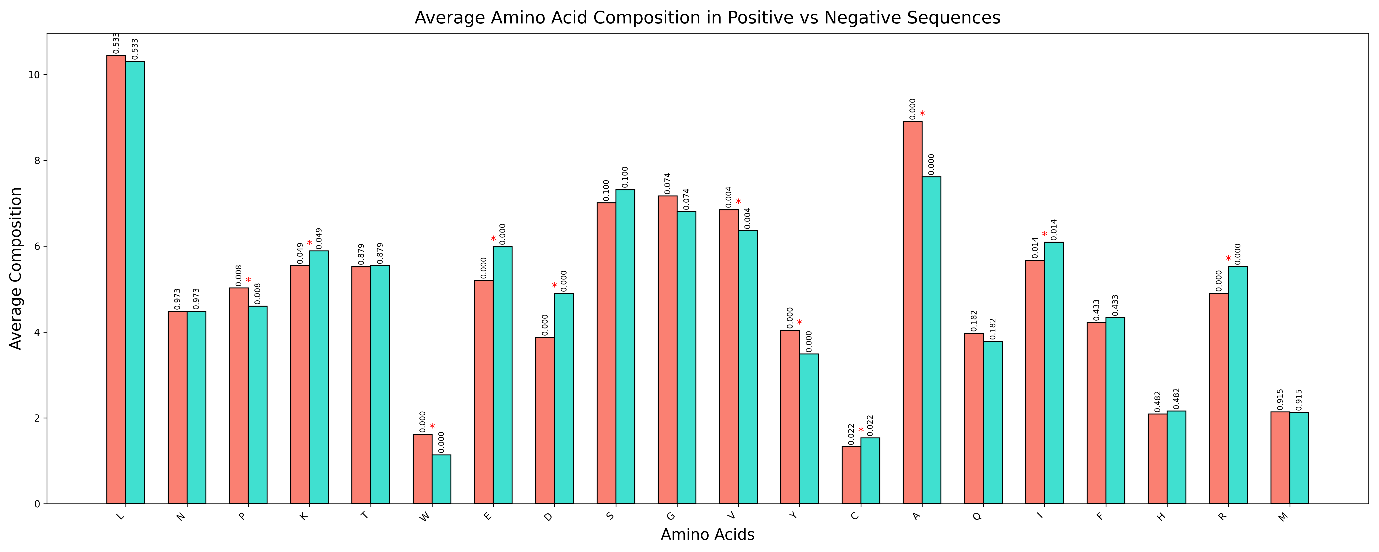

**Figure SF4:** Bar plot of the peptides' average single amino acid composition in the IL-2 inducers and non-inducers of the alternate dataset 2. The adjusted p-values are shown above the bars.

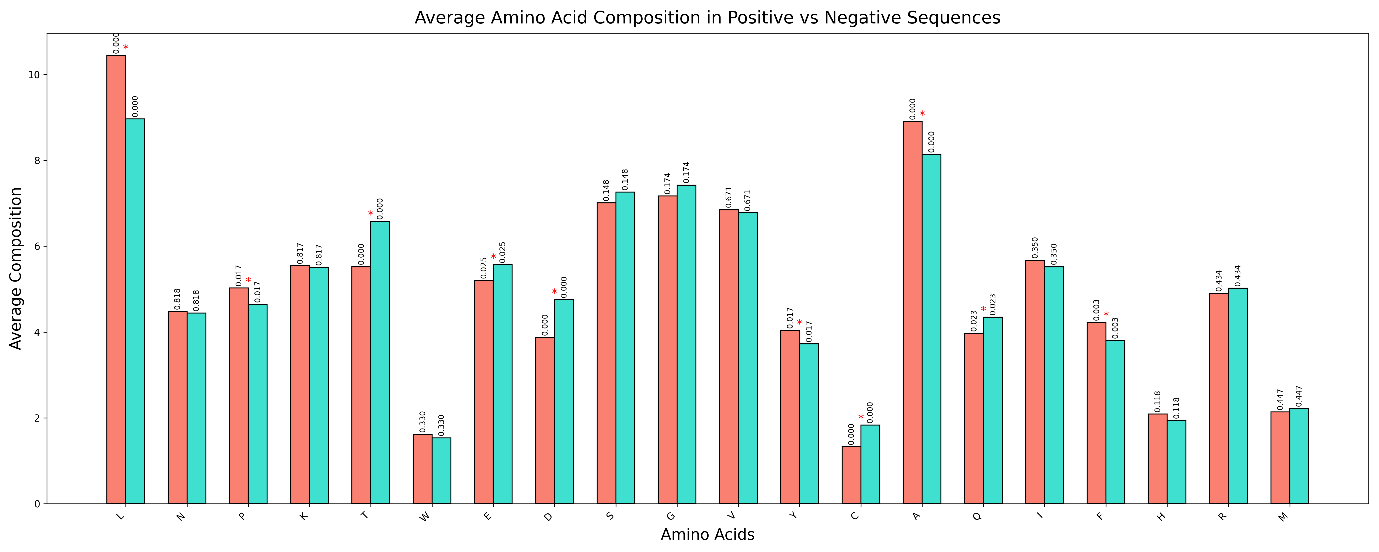

**Figure SF5:** Two-sample logo displaying the positional conservation of amino acid for the alternate dataset 1.

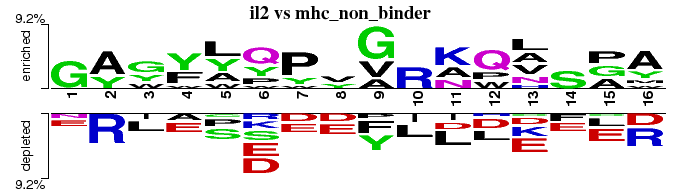

**Figure SF6:** Two-sample logo displaying the positional conservation of amino acid for the alternate dataset 2.

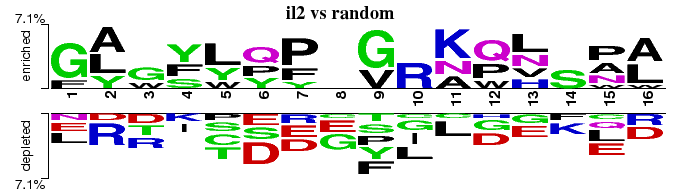
